# Molecular characterization of pathogen-induced SlCBL1-2, a calcium sensor, reveals the structural basis of calcium sensing in tomato

**DOI:** 10.64898/2026.10.07.757223

**Authors:** Sowrirajan Padmavathy Sowmiyarithika, Madhuravanam Cherukandy Amrutha, Muthukumar Bala Sundar, Mehanathan Muthamilarasan, Narayanasamy Sasirekha, Ponnuraj Karthe, Nagarathnam Radhakrishnan

## Abstract

Calcium (Ca^2+^) is a universal second messenger that orchestrates plant growth, development, and stress responses. Plant calcineurin B-like (CBL) proteins sense and relay Ca^2+^ signals in response to pathogens to activate downstream defense signalling pathways. Early blight disease, caused by the necrotrophic fungal pathogen *Alternaria solani*, affects tomato crops and severely limits their productivity. In the present study, we found that the tolerant tomato cultivar (Arka Rakshak) restricts *A. solani* infection and disease progression compared to the susceptible cultivar (PKM-1). Comparative transcriptome and qRT-PCR analysis of *A. solani*-infected tomato leaves showed that *SlCBL1-2* (Solyc08g007160.3), a Ca^2+^ sensor, is up-regulated in the tolerant cultivar, both relative to its own uninfected control and the infected susceptible cultivar, whereas it was not significantly induced in the uninfected susceptible cultivar. Protein sequence analysis indicates that SlCBL1-2 contains four EF-hand motifs, an N-terminal N-myristoylation/S-acylation module and a C-terminal FPSF motif, and it is predicted to interact with several CBL-interacting serine/threonine protein kinases (CIPKs). However, the evolutionary analysis, conformational transitions upon binding to Ca^2+^, and the Ca^2+^-binding affinity of SlCBL1-2 have not been previously characterized. Molecular Dynamics (MD) simulation of the AlphaFold structure of SlCBL1-2 indicates a well-folded structure and binding of two Ca^2+^ ions at EF-hands 3 and 4. Further, far-UV circular dichroism and tryptophan fluorescence spectroscopy of recombinant SlCBL1-2 indicate a natively folded structure in the absence of Ca^2+^ and undergo minimal secondary and tertiary structural changes upon Ca^2+^-binding, in contrast to several Ca^2+^-binding proteins. Of note, SlCBL1-2 binds Ca^2+^ with apparent K_D_ values of 526 ± 113 µM and 28.7 ± 9.47 µM, as evidenced by isothermal titration calorimetry. Together, our study provides an experimental description of the Ca^2+^-binding properties of tomato CBL protein and defines the molecular basis by which pathogen-induced SlCBL1-2 may act as a Ca^2+^ sensor in defense signalling.

## Introduction

Calcium (Ca^2+^) serves as a crucial second messenger in plants, and therefore, any changes in the cytosolic [Ca^2+^] levels activate the first line of defense responses in response to an external stimulus (Wang et al., 2023). When a plant encounters pathogen-associated molecular patterns (PAMPs), such as fungal chitin, pattern recognition receptors (PRRs) at the plasma membrane are activated and trigger a rapid increase in cytoplasmic Ca^2+^ ([Ca^2+^] _cyt_). The rise in [Ca^2+^] _cyt_ levels is mainly contributed by the intracellular Ca^2+^ stores such as the endoplasmic reticulum, vacuoles, chloroplasts, and mitochondria, and by the extracellular spaces. Various channels such as cyclic nucleotide-gated channels (CNGCs), glutamate receptor-like channels (GLRs), and two-pore channels (TPCs) elevate Ca^2+^ levels, giving rise to distinct “Ca^2+^ signatures” characterised by their frequency, duration, and amplitude depending upon the nature of the stimuli (Edel et al., 2017). The sudden rise in the intracellular Ca^2+^ levels is sensed and decoded by various Ca^2+^ sensors, leading to the activation of molecular cascades involved in defense responses (Vadassery & Oelmüller, 2009). Among the Ca^2+^ sensors, the CBL family has gained significant attention because of its specific role in decoding Ca²⁺ signals and kinase mediated downstream signalling through CBL-CIPK complex and adaptation to stress.

Plant CBL proteins were first identified in *Arabidopsis thaliana* (Liu & Zhu, 1998) and later characterized in other plant species including rice (Gu et al., 2008), chickpea (Poddar et al., 2022), pepper (Ma et al., 2019), turnip (Yin et al., 2017), brinjal (Li et al., 2016) and wild sugarcane (Feng et al., 2021). Plant CBL genes have been shown to play a pivotal role in responding to abiotic stress conditions such as drought (Zhu et al., 2023), salinity (Yang et al., 2019), flooding (Ye et al., 2018), and low temperature (Liu et al., 2024). Of note, SlCBL10 has been shown to have contrasting functions by acting as both a positive and negative regulator of salt tolerance in shoot and root, respectively (Estrada et al., 2023). In response to biotic stress such as fungal infection (*Rhizoctonia solani*) in potato, CBL4 has been shown to interact with CIPK2 and activate the downstream defense signalling against stem canker diseases (Yang et al., 2023). In the tobacco plant (*Nicotiana benthamiana*), overexpression of *CBLs* promotes disease resistance against bacterial wilt caused by *Ralstonia solanacearum* (Su et al., 2020).

Tomato (*Solanum lycopersicum* Linn.), a widely cultivated food crop worldwide, is affected by various fungal diseases (Panthee & Chen, 2010). Among the fungal diseases, early blight of tomato caused by *Alternaria solani* (Ellis and Martin) affects the whole plant, leading to defoliation and crop loss up to 80% (Thinakaran et al., 2020). Previous studies have reported the presence of 11 CBL-like proteins in tomato, and among them, SlCBL3-2 has been shown to negatively regulate under drought stress conditions (Tafreshi et al., 2021). However, the expression profile of CBL-like proteins under *A. solani* infection, the molecular aspects of Ca^2+^-binding responses, and the mechanism of CBL activating the downstream defense responses have not been previously characterized.

In our study, we showed that the tolerant tomato cultivar shows restricted *A. solani* colonisation and disease progression to early blight diseases. Transcriptomic analysis indicates that SlCBL1-2 is upregulated in the tolerant tomato cultivar compared to the susceptible cultivar infected with *A. solani.* Our evolutionary analysis indicates that CBL1 is a well-conserved protein among the Solanaceae members, however, it shows diversity among monocot and dicot plants. Molecular Dynamics (MD) simulation of SlCBL1-2 indicates a well-folded structure and binding of two Ca^2+^ ions at EF-hands 3 and 4. Consistent with this, far-UV circular dichroism and tryptophan fluorescence spectroscopy of purified SlCBL1-2 indicate a natively folded structure in the absence of Ca^2+^ and undergo minor secondary and tertiary structural changes upon Ca^2+^-binding. Of note, SlCBL1-2 binds Ca^2+^ with apparent K_D_ values of 526 ± 113 µM and 28.7 ± 9.47 µM, evident from Isothermal titration calorimetry. Our present study reports the Ca^2+^-binding properties of SlCBL1-2, integrating a multidisciplinary approach to understand the physiological function of SlCBL1-2 in tomato plants, which could serve as a potential target for improving tomato’s resistance against *A. solani* infection.

## Results and Discussion

### Tolerant tomato cultivar exhibits reduced *A. solani* colonization

In our study, we have infected the tolerant (Arka Rakshak) and susceptible (PKM-1) cultivars of tomato plants with the fungal pathogen, *A. solani,* to compare the disease progression/severity and to elucidate the molecular mechanism determining the tolerance (Adhikari et al., 2017). We used the double staining method combining Fluorescein Isothiocyanate-Conjugated Wheat Germ Agglutinin (FITC-WGA) and Propidium iodide (PI), which stain fungal and plant cell walls, respectively. Confocal laser scanning microscopy analysis at 8 days post-infection (dpi) reveals that the fungal pathogen penetrates the leaf tissue, colonizes, and spreads aggressively upon infection, leading to increased disease progression in the host plant (both tolerant and susceptible tomato cultivars) compared to their controls (**Fig. 1A**). Comparing the micrographs, it is evident that the *A. solani -*infected susceptible cultivar at 8 dpi revealed an increase in disease progression than the *A. solani -*infected tolerant cultivar. Further, the fluorescence intensity of WGA - FITC in the tolerant variety is significantly reduced, indicating restricted fungal colonisation, which could be due to the activation of host defense signalling responses (**Fig. 1B**).

**Fig. 1.**
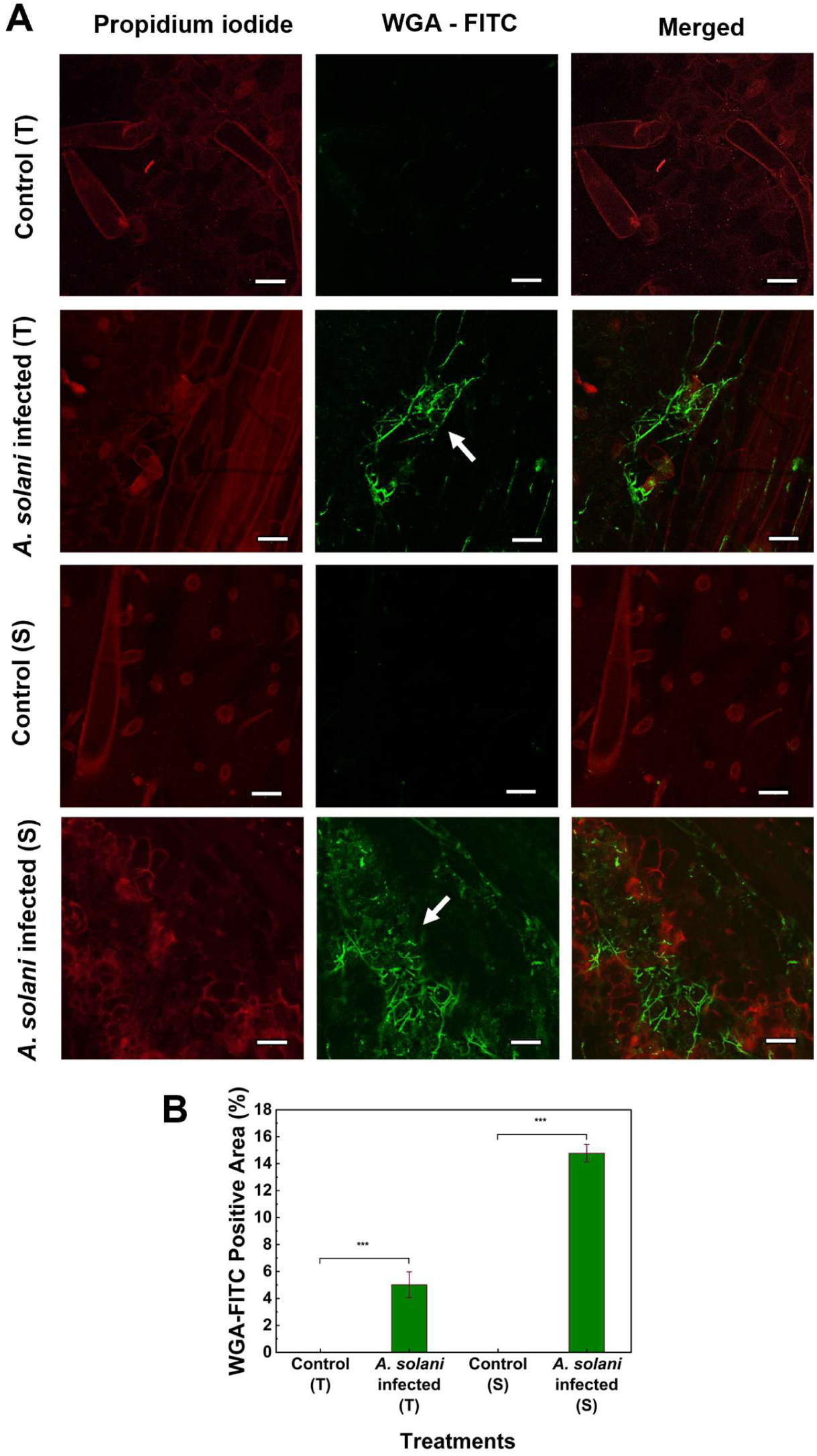
Restricted *A. solani* colonisation in the tolerant tomato cultivar. **A.** Confocal micrographs of tomato leaves of the tolerant cultivar (Arka Rakshak) and the susceptible cultivar (PKM-1) at 8 dpi, double-stained with FITC-WGA (fungus) and propidium iodide (plant cell wall). White arrows indicate fungal hyphae; colonisation is more extensive in the susceptible cultivar. Scale bar, 10 µm.**B.** Graph depicting the WGA- FITC positive area for both cultivars and treatments. *, **, and *** indicate significant differences between control and treated samples (i.e., p < 0.05; p <0.01 and p < 0.001, respectively) calculated using two-way ANOVA followed by Tukey’s test.

### Cultivar-specific transcriptional responses to *A. solani* infection

To further determine differences in gene expression levels, we performed comparative transcriptomic analysis between control and *A. solani*-infected tomato plants in tolerant and susceptible cultivars, as well as between *A. solani*-infected tolerant and *A. solani*-infected susceptible cultivars. For visualising the expression patterns of differentially expressed genes (DEGs), a hierarchical clustering heatmap was generated. The heatmap revealed a clear separation in expression profiles between the tolerant and susceptible cultivars, as well as between the infected and control conditions within each cultivar (**Fig. 2A**). Solyc08g007160.3 was designated as *SlCBL1-2* following the nomenclature of Tafreshi et al. (2021). In volcano plots, we observed that *SlCBL1-2* (Solyc08g007160.3) showed significant upregulation in the tolerant cultivar infected with *A. solani* (**Fig. 2B**) and in the *A. solani* infected tolerant vs *A. solani* infected susceptible cultivar (**Fig. 2F**). However, the gene was not significantly differentially expressed in the control vs *A. solani* infected susceptible cultivar (**Fig. 2D**). A significant differential gene expression was found to be more widely spread in the MA plot, which indicates the extensive transcriptional reprogramming induced by *A. solani*. The gene expression of *SlCBL1-2* was relatively lower in the susceptible cultivar, which might be due to the delayed defense response (**Fig. 2E**). In contrast, *SlCBL1-2* was upregulated in both control vs *A. solani* infected tolerant (**Fig. 2C**) and *A. solani* infected tolerant vs *A. solani* infected susceptible cultivar (**Fig. 2G**). Critically, *SlCBL1-2* (Solyc08g007160.3) was highlighted in MA plots, positioned above the baseline in the upregulated region. The venn diagram shows the genes that were upregulated (**Fig. 2H**) and downregulated (**Fig. 2I**) during the transcriptome analysis between control and *A. solani* infected tomato plants in tolerant and susceptible cultivar as well between *A. solani* infected tolerant vs *A. solani* infected susceptible cultivar.

**Fig. 2.**
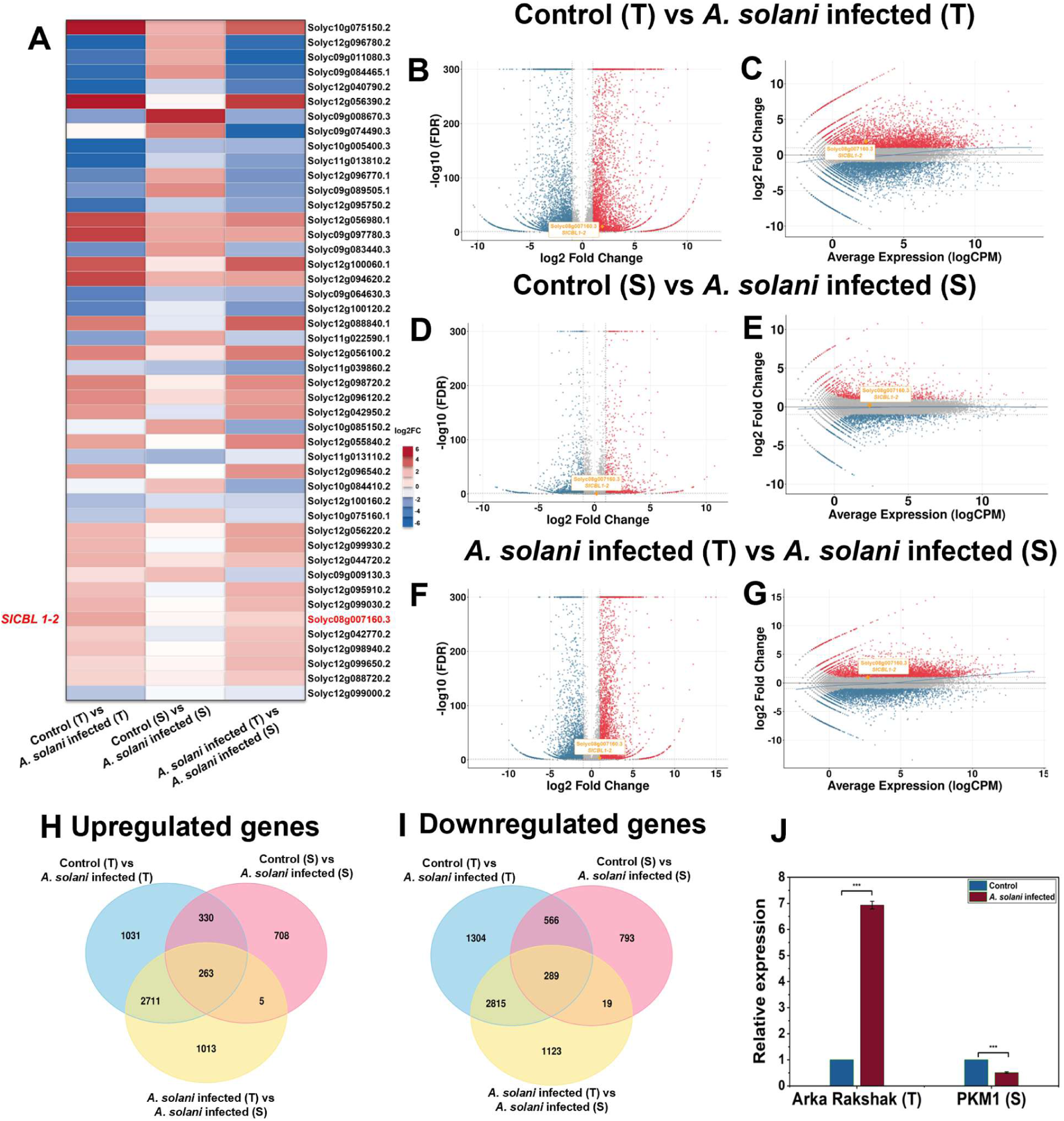
Transcriptome analysis of tolerant and susceptible tomato cultivars infected with *A. solani*. **A.** Heatmap representation of *SlCBL1-2* gene expression in Control (Tolerant), *A. solani-*infected (Tolerant), Control (Susceptible), and *A. solani-*infected (Susceptible). The heatmap color scale ranged from -2 to +2 (log2-normalised expression), where red indicates high relative expression, and blue indicates low relative expression. Volcano plots of differentially expressed genes identified between Control (Tolerant) vs *A. solani-*infected (Tolerant)(**B**), Control (Susceptible) vs *A. solani-*infected (Susceptible)(**D**), and *A. solani-*infected (Tolerant) vs *A. solani-*infected (Susceptible)(**F**). MA plot analysis of differentially expressed genes identified between Control (Tolerant) vs *A. solani-*infected (Tolerant)(**C**), Control (Susceptible) vs *A. solani-*infected (Susceptible)(**E**), and *A. solani-*infected (Tolerant) vs *A. solani-*infected (Susceptible)(**G**). Venn diagram of genes that were upregulated (**H**) and downregulated (**I**) between Control (Tolerant) vs *A. solani-*infected (Tolerant), Control (Susceptible) vs *A. solani-*infected (Susceptible), and *A. solani*-infected (Tolerant) vs *A. solani-*infected (Susceptible). **J.** Validation of the *SlCBL1-2* gene using qRT-PCR in control and *A. solani-*infected leaves in tolerant and susceptible tomato cultivars. Statistical significance was performed using two-way ANOVA followed by Tukey’s test and defined as * (p < 0.05), **(p <0.01), and *** (p < 0.001).

### Upregulation of *SlCBL1-2* in *Alternaria solani-*infected tolerant tomato cultivar

The transcriptome result was further validated by quantitative real-time PCR (qRT-PCR) on control and *A. solani*-infected leaves of the tolerant and susceptible cultivars at 8 dpi, using Tubulin as the reference gene. *SlCBL1-2* transcript levels were strongly induced in the infected tolerant cultivar relative to its own uninfected control, whereas in the susceptible cultivar, the transcript level in infected leaves was lower than in its control (**Fig. 2J**). The qRT-PCR results were consistent with the transcriptome results, which supports the differential regulation of *SlCBL1-2* in response to *A. solani* infection in tolerant and susceptible cultivars. This result implies that *SlCBL1-2* shows a stronger transcriptional response in the tolerant cultivar compared to the susceptible cultivar.

This cultivar-specific expression pattern of *SlCBL1-2* from this analysis strongly indicates the Ca^2+^-mediated defense signalling cascades in the tolerant tomato cultivar during *A. solani* infection. Similar studies from other laboratories have shown that the *SaCBL1-like* gene, which was upregulated only in *Solanum acaule* (wild potato), confers freezing tolerance and was found to interact with SaCIPK3–1(Chen et al., 2022). Also, in soybean (*Glycine max*), GmCBL1 expression is increased in response to saline and alkaline stress (Jiao et al., 2024). Of note, in *Arabidopsis thaliana*, *CBL5* has been reported as essential for seed germination, as the mutant inhibits germination due to salt stress and is reported to interact with CIPK8 or CIPK24 (Xie et al., 2024).

### CBL1-2, a Ca^2+^ sensor of *Solanum lycopersicum,* might be involved in pathogen-induced Ca^2+^ signalling mechanism

Primary sequence analysis indicates that SlCBL1-2 contains 215 amino acid residues with a theoretical molecular weight of 24.65 kDa. Grand Average of Hydropathicity (GRAVY) score of -0.228 indicates that SlCBL1-2 is hydrophilic in nature and a soluble protein. STRING analysis predict that SlCBL1-2 interacts with several CBL-interacting serine/threonine protein kinases (CIPKs) such as CIPK9 (A0A3Q7GL89), CIPK23 (A0A3Q7FGM7) and CIPK24 (SOS2) (**Supplementary Figure 1**). CIPKs are known to phosphorylate CBL in a CBL-CIPK complex and results in the downstream signal transduction of target proteins, such as K+ transporter (AKT1), high-affinity K+ transporter 5 (HAK5) and ion channels including Slow Anion Channel-Associated 1 (SLAC1) and S-type anion channels, and regulates transcription of stress-associated genes (Chen et al., 2024). Additionally, SlCBL1-2 is predicted to interact with AKT1 (a potassium channel) and TPC1 (a two-pore Ca^2+^channel). Studies in other plants, such as *A. thaliana* have shown that CBL1 and 9 interact with CIPK23, and this complex activates AKT1 for regulating the K+ influx across the plasma membrane under potassium deficient condition. (Cheong et al., 2007). These analyses indicate that upregulation of SlCBL1-2 in tolerant tomato cultivars might also initiate a similar cascade of Ca^2+^-mediated signal transduction events following *A. solani* infection to induce defense-related genes.

Our analysis further indicates that SlCBL1-2 contains an N-myristoylation site (MGCXXSK) at the N-terminus, which is also conserved among Solanaceae members. Adjacent to the N-myristoylation site (Glycine), a cysteine residue is located, which is predicted as a putative S-acylation (palmitoylation) site. Both the N-myristoylation and S-acylation sites in SlCBL1-2 suggest the potential dual lipid modification in the protein, necessary for membrane localisation. Of note, N-myristoylation in human neuronal calcium sensor-1 (NCS-1) is crucial for insertion into the plasma membrane (Baksheeva et al., 2020). In addition, SlCBL1-2 also have a well-conserved FPSF motif which harbours the phosphorylation site at the C-terminus, which is known to be phosphorylated by CBL-interacting protein kinase (CIPK) in other plants. (**Fig. 3B**). However, the functional significance of these motifs is not understood in SlCBL1-2.

**Fig 3.**
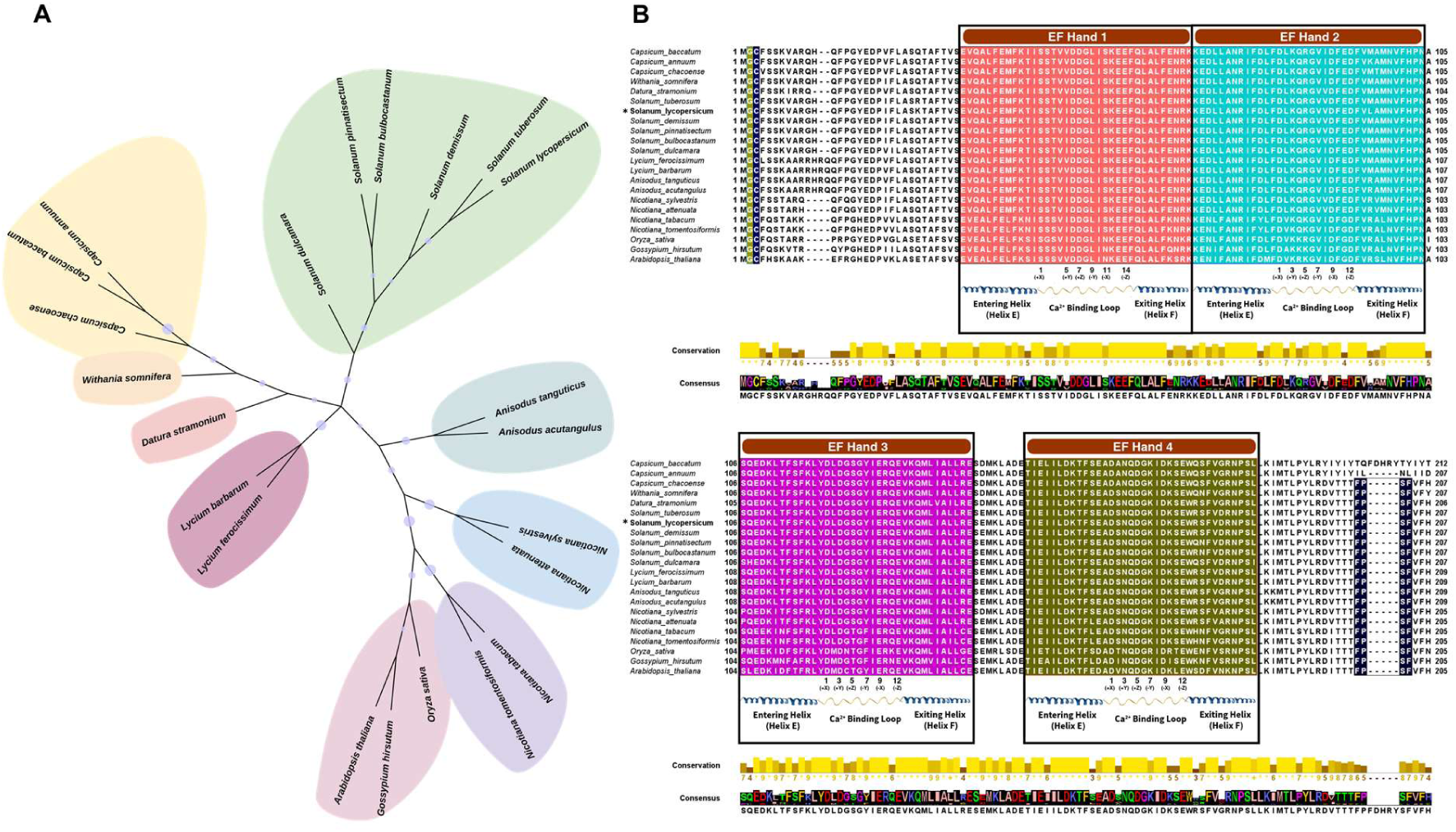
Evolutionary and sequence analysis of CBL1 proteins. **A.** The phylogenetic analysis of CBL1 among Solanaceae members using MEGA11 software (1000 bootstrap) with *Arabidopsis thaliana*, *Gossypium hirsutum,* and *Oryza sativa* as outgroups. **B.** Multiple sequence alignment of CBL1 across the Solanaceae family members with *Arabidopsis thaliana*, *Gossypium hirsutum,* and *Oryza sativa* as reference. The amino acids corresponding to EF-hand 1(highlighted in red), EF-hand 2 (highlighted in blue), EF-hand 3 (highlighted in pink), and EF-hand 4 (highlighted in brown). The N- myristoylation (highlighted in yellow) and palmitoylation site (highlighted in blue) and the FPSF site (highlighted in black) are well conserved across the Solanaceae members.

### Evolutionary relationship of CBL Proteins in Solanaceae family members

To assess the evolutionary relationship of SlCBL1-2 among the Solanaceae family members, a phylogenetic tree was constructed using maximum likelihood (ML) method. We have taken the full-length amino acid sequence of CBL1 proteins belonging to Solanaceae members for multiple sequence alignment and phylogenetic tree construction. The CBL1 protein sequences of *Arabidopsis thaliana* (thale cress), *Gossypium hirsutum* (cotton) and *Oryza sativa* (rice) were used as the outgroup. Our analysis indicates that CBL is a well-conserved protein among the Solanaceae members. In addition, the CBL1 proteins are widely distributed across monocot and dicot plants, while retaining sequence conservation, thereby indicating the evolutionary conservation of their roles in Ca^2+^-mediated signalling processes. **(Fig. 3A).** Further our analysis indicates that SlCBL1-2 protein shows high sequence conservation with CBL1 of *Solanum tuberosum* and shares 99% sequence identity. The CBL proteins in plants share sequence similarity with calcineurin B subunit 1(CNB1) in yeast and neuronal calcium sensor 1(NCS1) in human (Mao et al., 2016; Weinl & Kudla, 2009). Multiple sequence alignment of neuronal calcium sensor 1 in *Homo* sapiens, calcineurin subunit B in *Saccharomyces cerevisiae*, AtCBL1 in *Arabidopsis thaliana* and SlCBL1-2 of *Solanum lycopersicum* reveals strong conservation in the EF-hand motifs (**Supplementary Figure 2**). In addition, amino acid residues involved in Ca^2+^ binding were found to be conserved in EF-hands 2, 3 and 4 in all the sequences. Unlike human (NCS1) and fungal (CNB1) Ca^2+^ sensors, the EF-hand 1 in plant CBL (AtCBL1 and SlCBL1-2) displayed divergence from the canonical EF-hand sequence by having a long Ca^2+^ coordination loop, indicating that the protein might have undergone evolutionary changes to accommodate plant Ca^2+^ signalling.

### Structural conservation of EF-hand loop residues in SlCBL1-2

Secondary structure analysis indicates that SlCBL1-2 contains four-EF-hand motifs, which are well-conserved among Solanaceae members, especially in the Ca^2+^-binding loop regions. Comparing the four EF-hands of SlCBL1-2 against the canonical EF-hand Ca^2+^-binding consensus sequence, Ca^2+^ is coordinated by 6 amino acid residues through the side/main chain oxygen atoms represented as +X, +Y, +Z, -Y, -X, and -Z, based on the pentagonal bipyramidal geometry (Gifford et al., 2007). Asp (D), Asn (N), Ser (S), or Glu (E) are the most common amino acids found in canonical EF-hands, as they have side chains that contain oxygen atoms. In addition, Gly (G) is highly conserved in the middle of the EF-hand loop to allow the loop to fold around the Ca^2+^. Interestingly, the -Z position is usually occupied by Glu (E) for bidentate Ca^2+^ coordination in canonical EF-hands.

In contrast to other Ca^2+^ sensors that contain 12 amino acid residues, the first EF-hand in plant CBLs contains 14 amino acid residues, as there is a two-amino-acid insertion between the +X and +Y positions (Hunpatin et al., 2023). Our analysis indicates that EF-hand 1 of SlCBL1-2 shows conservation at Ser (S) (+X), Asp (D) (+Z), and Glu (E) (-Z) positions similar to EF-1 of *A. thaliana* **(Table 1)**. However, the EF-loop has Ile (I) and Leu (L) at +Y and −Y positions and lacks side-chain oxygen atoms essential for Ca^2+^ coordination. Therefore, EF-hand 1 of SlCBL1-2 is a non-canonical EF-hand and might serves as a putative Ca^2+^ binding site.

**Table 1.** Comparison of amino acid residues at the canonical Ca²⁺-coordinating positions in EF-hand 1 of SlCBL1-2 with the AtCBL1 EF-hand motif.

| Residue Number | 1 | 2 | 3 | 4 | 5 | 6 | 7 | 8 | 9 | 10 | 11 | 12 | 13 | 14 |
| --- | --- | --- | --- | --- | --- | --- | --- | --- | --- | --- | --- | --- | --- | --- |
| AtCBL1 EF-Hand 1 | +X<br>(S) |  |  |  | +Y<br>(I) |  | +Z<br>(D) | (G) | -Y<br>(L) |  | -X<br>(N) |  |  | -Z<br>(E) |
| SICBL1-2 EF-Hand 1 | S | S | T | V | I | D | D | G | L | I | S | K | E | E |

All the other EF-hands (i.e., 2, 3 and 4) have Asp (D) in the +X position, which is essential for Ca^2+^ coordination (**Table 2**). The +Y position is conserved only in EF-hands 3 (Asp (D)) and 4 (Asn (N)), and Lys (K) occupies that position in EF-hand 2. Additionally, Arg (R) and Ser (S), respectively, were found at (+Z) in EF-hands 2 and 3. Of note, Gly (G), which plays a pivotal role in conferring unusual main-chain conformation of the EF-loop, is conserved among all the 4 EF-hands of SlCBL1-2. Similarly, the (-Z) position, which is usually occupied by Glu (E), is highly conserved among all EF-hands.

**Table 2.** Comparison of amino acid residues at the canonical Ca²⁺-coordinating positions across EF-hands 2–4 of SlCBL1-2 with the canonical EF-hand motif.

| Residue Number | 1 | 2 | 3 | 4 | 5 | 6 | 7 | 8 | 9 | 10 | 11 | 12 |
| --- | --- | --- | --- | --- | --- | --- | --- | --- | --- | --- | --- | --- |
| Canonical EF-Hand | +X<br>(D) |  | +Y<br>(D/N) |  | +Z<br>(D/N/<br>S) | (G) | -Y |  | -X |  |  | -Z<br>(E) |
| SICBL1-2 EF-Hand 2 | D | L | K | Q | R | G | V | V | D | F | E | D |
| SICBL1-2 EF-Hand 3 | D | L | D | G | S | G | Y | I | E | R | Q | E |
| SICBL1-2 EF-Hand 4 | D | S | N | Q | D | G | K | I | D | K | S | E |

### Molecular Dynamics (MD) simulation studies of SlCBL1-2

Next, we sought to study the impact of Ca^2+^-binding on protein folding, stability, Ca^2+^-induced conformational changes, and loop fluctuations in SlCBL1-2. We used the AlphaFold structure of SlCBL1-2 for the MD simulation since the experimentally derived structure is not currently available. We carried out MD Simulation studies using GROMACS 5.0 for the SlCBL1-2 protein with Ca^2+^ and without Ca^2+^ for 100 ns **(Figs. 4A-E)**. For the Ca^2+^ bound structure, we initially performed Ca^2+^ binding site prediction and docking using MIB2 server, which predicted EF-hands 3 and 4 to be Ca^2+^ binding regions. To further validate the docking predictions, Ca^2+^ ions were added in the neutralisation step prior to running MD simulation. Interestingly, similar to the MIB2 prediction, the Ca^2+^ ions remained stably sequestered within the Ca^2+^ -binding pockets of EF-hands 3 and 4 but not in the EF-hand 1 and 2 throughout the simulation trajectory. Furthermore, the root mean square deviation (RMSD), which indicates the stability of the SlCBL1-2 during the simulation time compared to its initial conformation was evaluated. Three independent MD replicates were performed and the RMSD and RMSF values from the three replicas were averaged. The average RMSD values of the apo and holo SlCBL1-2 protein were found to be 0.66 and 0.51 nm, respectively indicating that SlCBL1-2 does not undergo major conformational changes in the presence/absence of Ca^2+^. The RMSD trajectory reveals that the SlCBL1-2 protein undergoes significant structural deviations in its apo form, reaching values as high as 0.78 nm. Conversely, the RMSD of Ca^2+^-bound SlCBL1-2 reached a plateau much earlier and maintained a significantly lower RMSD, stabilizing between 0.45 nm and 0.55 nm, indicating that Ca^2+^-binding induces a more compact and stable conformation in SlCBL1-2 **(Fig. 4F)**.

**Fig. 4.**
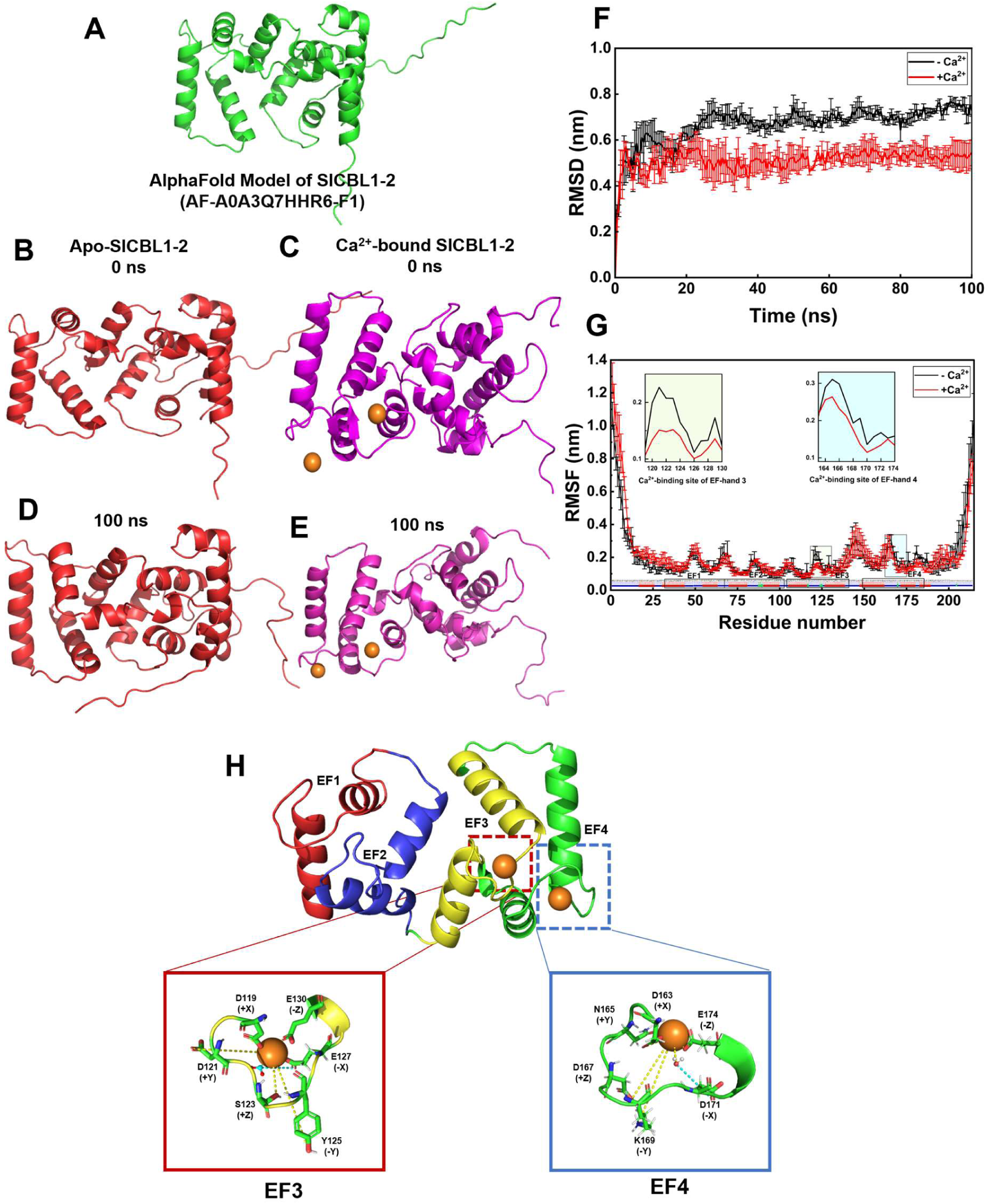
Molecular dynamics simulation of apo and Ca²⁺-bound SlCBL1-2 reveals Ca^2+^-mediated structural stabilisation. **A.** AlphaFold structure of SlCBL1-2 protein was subjected to molecular dynamics simulation for 100 ns in the presence and absence of Ca^2+^. Structure of apo-SlCBL1-2 protein at 0 ns **(B)** and 100 ns **(C).** Structure of Ca^2+^-bound SlCBL1-2 protein at 0 ns **(D)** and 100 ns **(E). (F)** Root Mean Square Deviation (RMSD) for apo (black) and Ca²⁺-bound (red) SlCBL1-2 during 100 ns MD simulation. **(G)** Root Mean Square Fluctuation (RMSF) of apo (black) and Ca²⁺-bound (red) SlCBL1-2. **(H)** Structural organisation of the four EF-hand motifs (EF1–EF4) in SlCBL1-2, with Ca²⁺ ions shown as orange spheres bound specifically to EF-hand 3 and EF-hand 4.

To interpret the conformational changes induced by Ca^2+^-binding at the EF-hand loop regions in SlCBL1-2, root mean square fluctuation (RMSF) was calculated. From the RMSF plot, we observe fluctuations in the EF-hand 3 loop (from Asp^119^ to Tyr^125^) between the apo and Ca^2+^-bound forms, with a difference of 0.05 nm. Additionally, a minimal fluctuation was observed between the Asp^148^ to Glu^161^ amino acid residues in the entering helix of the EF-hand 4, indicating a slight conformational change due to Ca^2+^-binding to the EF-hands **(Fig. 4G)**. MD simulation studies further indicated the Ca^2+^ coordination sites at EF-hands 3 and 4 **(Fig. 4H)**. In EF-hand 3, the amino acids in the loop region, Asp^119^(+X), Asp^121^(+Y), Ser^123^(+Z), Tyr^125^(-Y), Glu^127^(-X) and Glu^130^(-Z) coordinate a Ca^2+^ ion. Similar sites in EF-hand 4 loop regions (i.e., Asp^163^(+X), Asn^165^(+Y), Asp^167^(+Z), Lys^169^(-Y), Asp^171^(-X) and Glu^174^ (-Z)) are involved in binding of Ca^2+^ion.

According to Bork et al., (2024), AtCBL1 has three Ca^2+^-binding sites, where EF-hand 1 could bind Ca^2+^, because of its non-canonical loop. Similarly, EF-hand 1 of SlCBL1-2 have a sequence-level variation within the non-canonical loop, which makes Ca^2+^-binding not feasible at EF-hand 1. However, we observe that EF-hand 2 also does not bind Ca^2+^, and EF-hands 3 and 4 function only as the Ca^2+^-binding sites in SlCBL1-2.

To study the biophysical properties and understand the Ca^2+^-binding responses of SlCBL1-2, we have subcloned the *SlCBL1-2* (648 bp; Uniport ID: A0A3Q7HHR6) gene into the bacterial expression vector and purified it using Ni-NTA affinity and gel-filtration chromatography (**Fig. 5A & 5C**). The protein (Mol. Wt: 28 kDa) was confirmed by western blotting using anti-His tag antibody (**Fig. 5B**), which corresponds to the molecular weight of SlCBL1-2 protein with His tag.

**Fig. 5.**
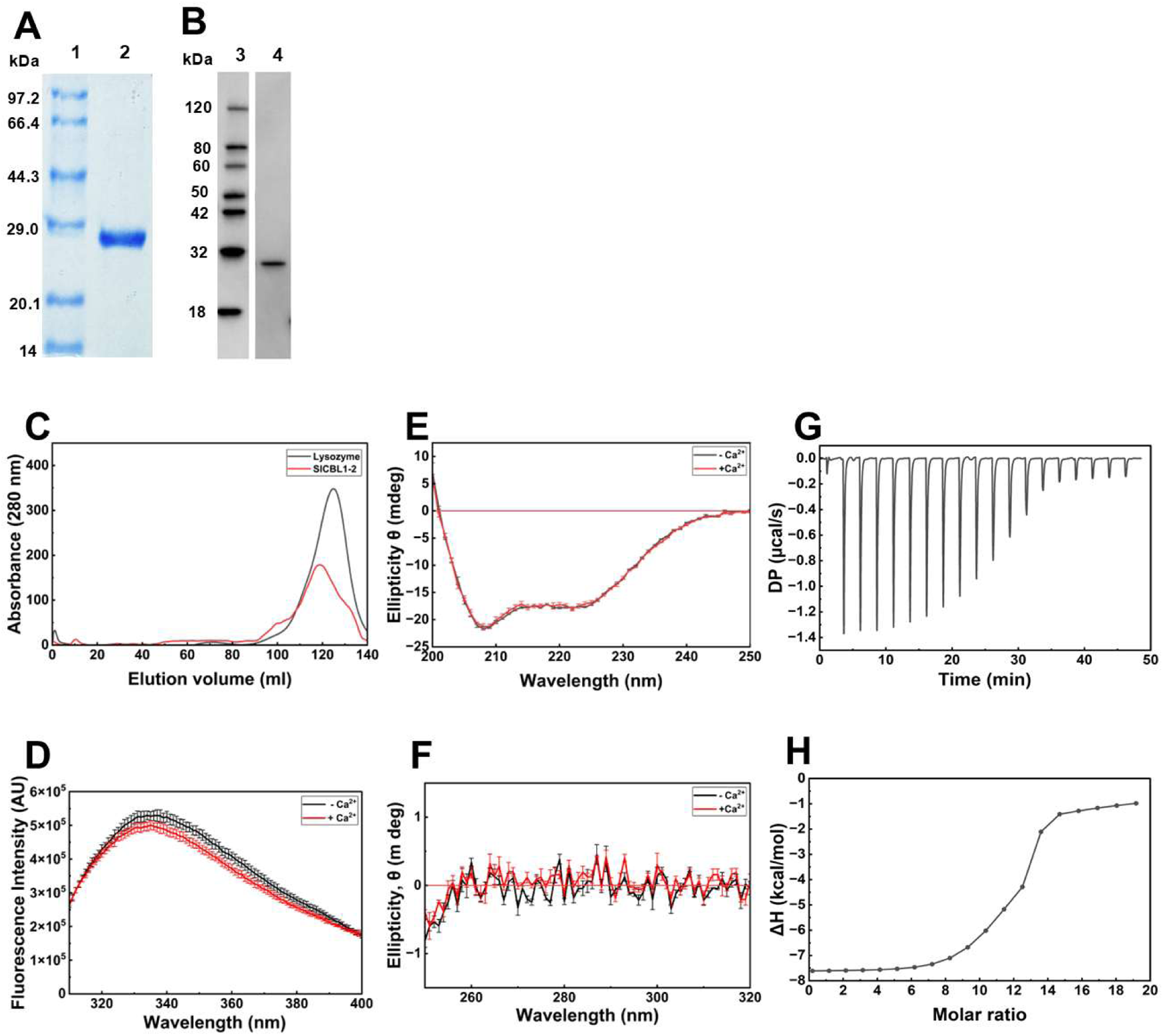
Structural changes of SlCBL1-2 upon Ca²⁺-binding. **A.** SlCBL1-2 protein purified using S75 Column on 12.5 % SDS PAGE Lane 1 – Low molecular Weight Protein Marker; Lane 2–SlCBL1-2 protein **B.** Western blot analysis of SlCBL1-2 using anti-His antibody with Lane 3 – Low molecular weight marker; Lane 4 – SlCBL1-2 protein. **C.** Gel Filtration chromatography using S75 column for lysozyme and SlCBL1-2 protein **C.** Far UV CD spectra were recorded for SlCBL1-2 protein (30 µM) in 20 mM Tris, 50 mM NaCl (pH 7.4) in the absence and presence of Ca^2+^(1 mM). **D.** Fluorescence emission spectra of SlCBL1-2 protein (2 µM) in 20 mM Tris, 50 mM NaCl (pH 7.4), with excitation at 295 nm and emission spectra recorded from 310 to 400 nm. The spectra were recorded for apo and holo protein (1 mM Ca^2+^). **E.** Near-UV CD spectra were recorded for SlCBL1-2 protein (50 µM) in 20 mM Tris, 50 mM NaCl (pH 7.4) in the absence and presence of Ca^2+^(1 mM). **F.** Far-UV CD spectra were recorded for SlCBL1-2 protein (50 µM) in 20 mM Tris, 50 mM NaCl (pH 7.4) in the absence and presence of Ca^2+^(1 mM). **G.** Raw ITC thermogram showing heat changes upon successive injections of 1 mM of CaCl₂ (2 µL per injection) into 10 µM SlCBL1-2 protein **H.** Binding isotherm showing Ca²⁺-binding to SlCBL1-2, fitted using two set of sites model.

### Ca^2+^-binding induces minor structural changes in SlCBL1-2

SlCBL1-2 contains a single Trp residue located at the exiting helix of the EF-hand 4 adjacent to the Ca^2+^-binding site (**Fig. 5D**). Therefore, we used fluorescence spectroscopy to monitor the conformational changes in SlCBL1-2 upon binding to Ca^2+^ using Trp as an intrinsic probe. SlCBL1-2 was excited at 295 nm, and the emission spectra were recorded from 310-400 nm. SlCBL1-2 shows maximum fluorescence emission (λmax) at 335 and 334 nm, in the presence and absence of Ca^2+^, respectively. The negligible shift in the emission maxima between Ca^2+^-free and bound forms suggests that the microenvironment around the Trp residue is not altered by Ca^2+^-binding to the EF-hand regions in SlCBL1-2.

We performed circular dichroism (CD) spectroscopy to detect the secondary structural changes of SlCBL1-2 in the presence and absence of Ca^2+^. CD spectra for apo-SlCBL1-2 in the far-UV region (200-250 nm) exhibit characteristic negative ellipticity peaks at 208 nm and 222 nm, indicative of α-helical regions corresponding to the EF-hands and a natively folded structure in the absence of Ca^2+^. Upon addition of Ca^2+^, no significant changes in the spectra were observed, consistent with MD simulation studies (**Fig. 5E**). These results indicate that Ca^2+^-binding did not induce any conformational changes/alter the secondary structure of the protein. Using the BeStSel online tool, we quantified the alpha-helical content of the apo and holo protein from the CD spectra. We observed that the alpha-helical structure of the protein was estimated to be 53% in the absence of Ca^2+^. In the presence of Ca^2+^, the alpha-helical content increased to 55% (Table S2). These results suggest that the SlCBL1-2 protein maintained its overall alpha-helical structure in the presence and absence of Ca^2+^.

Further near-UV (250 – 300 nm) CD spectroscopy was done to monitor the tertiary structural changes in SlCBL1-2. The near-UV CD signals are contributed by the aromatic amino acid residues, with phenylalanine contributing in the 255–270 nm region, tyrosine around 275–282 nm, and tryptophan in the region near 285–305 nm. SlCBL1-2 has 1 Trp, 4 Tyr, and 20 Phe residues. The near-UV spectrum of apo SlCBL1-2 shows minimal ellipticity near the baseline, consistent with tight packing of aromatic side chains. Ca^2+^-binding resulted in a slight increase in ellipticity between 255–295 nm, indicative of minor local conformational changes, decreased mobility of the aromatic residues, and a subtle rigidity of the tertiary structural organization (**Fig. 5F**). However, the absence of distinct peaks around the Trp residue (∼285 – 305 nm) suggests that no significant exposure of aromatic amino acid residues occurs upon Ca²⁺-binding to the protein, consistent with the Trp fluorescence measurement. A minimal fluctuation around the Phe residue (∼255–270 nm) and the weak signals near the Tyr (∼275–282 nm) suggest that the tertiary structure of the protein remains stable irrespective of Ca^2+^-binding.

Together, these results suggest that Ca^2+^-binding stabilizes the overall structure with minimal secondary and tertiary structural changes. However, few EF-hand proteins in plants, including calmodulin-like protein 15 (CML15) and calmodulin-like protein 16 (CML16) in *Arabidopsis thaliana* are predominantly alpha-helical proteins, as evident from far-UV CD spectroscopy. Upon addition of Ca^2+^, CML15 presented a decrease in the helical content, whereas CML16 displayed an increase in the helical content (Ogunrinde et al., 2017). Similarly, calcium-dependent protein kinase 1 (CDPK1) from *Cicer arietinum* indicated that the protein increases its helical content upon the addition of Ca^2+^(Dixit & Jayabaskaran, 2015). Unlike other plant Ca^2+^ sensors, SlCBL1-2 is stable both in Ca^2+^-bound and free forms.

### Thermodynamic characterization of Ca^2+^ binding to SlCBL1-2

To determine the Ca^2+^-binding affinity of SlCBL1-2 and the associated binding parameters, Isothermal Titration Calorimetry (ITC) was performed. Ca^2+^-binding to SlCBL1-2 is an exothermic reaction, as evident from the release of heat during each injection (**Fig. 5G & 5H**). The binding isotherm was fitted to two sets of binding sites with the K_D_ values of 526 and 28.7 µM, respectively **(Table 3)**. Our study indicates that Ca^2+^-binding to SlCBL1-2 is a thermodynamically favourable reaction, as evidenced by the negative Gibbs free energy. The negative enthalpy observed in both sites was likely due to the favourable electrostatic interaction between the Ca^2+^ and the conserved acidic amino acid residues within the EF-hand loops, and the entropy indicates a more structurally ordered state. Our findings are consistent with fluorescence and CD spectroscopic analyses, as SlCBL1-2 exists as a natively folded alpha-helical protein and undergoes minor secondary and tertiary structural changes upon Ca^2+^ binding. These results indicate that despite being a folded protein, SlCBL1-2 binds Ca^2+^ with low-to-moderate affinity. The derived Ca^2+^-binding affinity values are comparable to other Ca^2+^ sensors in plants such as calmodulin-like protein 14 (CML14) from *Arabidopsis thaliana*, where the protein binds to a single Ca^2+^-binding site with a dissociation constant of 12 µM. Similarly, Ogunrinde et al., (2017) reported that CML15 binds two Ca^2+^ with K_D_ of 0.22 and 1.23µM and CML16 has three functional Ca^2+^-binding sites with 0.03, 0.21, and 3.76µM Ca^2+^-binding affinities. These results highlight the diversity in plant Ca^2+^ sensors and their ability to decode distinct Ca²⁺ signatures (Vallone et al., 2016). Collectively, our results indicate that SlCBL1-2 might function as a Ca^2+^ sensor in tomato plants, decode Ca^2+^ signals during pathogen attack and be involved in downstream signalling pathways during biotic stress.

**Table 3:**
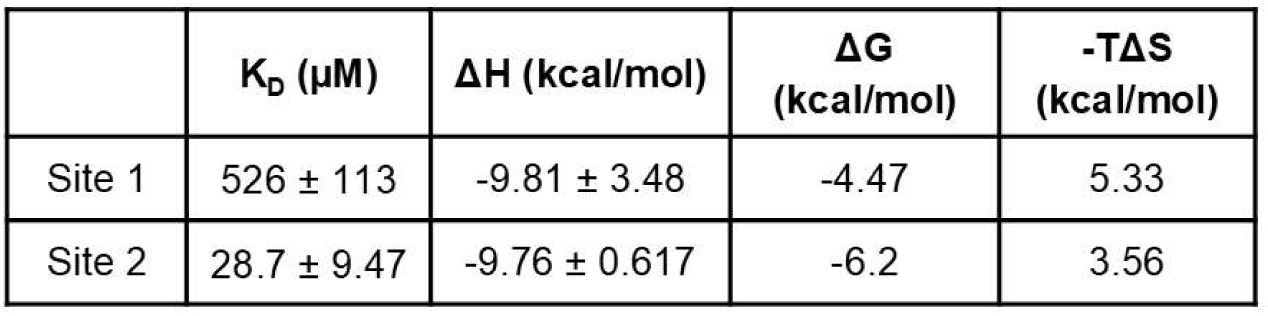
Thermodynamic parameters of Ca^2+^-binding to SICBL1-2.

In our present study, we found that the tolerant cultivar of tomato showed reduced fungal ramification compared to the susceptible cultivar. Comparative transcriptomic analysis indicates that the SlCBL1-2 gene is upregulated upon *A. solani* infection, indicating the potential involvement of Ca^2+^-signalling systems in tomato plant defense responses. Our studies report the molecular characterization of SlCBL1-2, a Ca^2+^ sensor, and indicate that SlCBL1-2 is a natively folded protein that doesn’t undergo major conformational changes upon binding to Ca^2+^. However, it binds to Ca^2+^ with low-to-moderate affinity binding sites and might be involved in the activation of CIPKs. Further experiments are warranted to study the effect of SlCBL1-2 and CIPK interactions, which would open many avenues in understanding tomato-*Alternaria* interaction, their physiological significance, and could be potentially targeted for improving plant resistance against *A. solani* infection.

## Materials and Methods

### Plant materials and pathogen infection

Seeds of *Solanum lycopersicum* cv. Arka Rakshak (tolerant) and cv. Periyakulam-1 (PKM-1) (susceptible) to early blight disease, were procured from the ICAR - Indian Institute of Horticultural Research (ICAR-IIHR) and Tamil Nadu Agricultural University (TNAU), India respectively. The tomato plants were grown in pots containing soil mixed with vermiculite under greenhouse conditions with 16 hours of daylight and 8 hours of darkness. The fungal pathogen, *Alternaria solani* (Accession No. 4632), was procured from ICAR-Indian Agricultural Research Institute (ICAR- IARI), New Delhi, India. For the preparation of *A. solani* suspension, the fungal pathogen was grown on corn meal agar medium for 7 days and the mycelium was scraped off from the surface. After twenty-five days, healthy tomato plants were infected with *A. solani* (2 × 10^6^ conidia mL^−1^) by foliar spraying, and the control plants were treated with water (Dey et al., 2019). The appearance of the first visible disease symptom was considered Day 0. All the experimental leaf samples were harvested at 8 dpi, and visualised under a confocal microscope as described by Irullapan *et al*., (2022). WGA-FITC and Propidium Iodide were used for staining the fungal culture and the leaf, respectively. The microscopy observations were performed under a multiphoton confocal microscope (Carl Zeiss, Germany) with excitation and emission bandpass of 495 and 515 nm for WGA-FITC and 535 nm and 617 nm for Propidium iodide.

### Sample preparation and Sequencing

The total RNA was extracted from the leaf samples of tolerant (Arka Rakshak) and susceptible (PKM-1) tomato cultivars under control (uninfected) and *A. solani*-infected conditions at 8 days post-inoculation (dpi). Three biological replicates were used for all treatment. Paired-end transcriptome sequencing libraries were prepared from the high-quality total RNA samples and Illumina sequencing was performed. The sequencing was carried out by Redcliffe Life Sciences Pvt. Ltd. (Noida, India). Pre-processing with fastp (v0.20.1) yielded high-quality filtered reads ranging from 93.79% to 98.32% of the total raw reads across all samples. High-quality filtered reads were aligned to the *Solanum lycopersicum* reference genome (SL4.0) using HISAT2 (v2.1.0), a fast and sensitive splice-aware alignment program that uses a graph-based alignment of reads to a population of genomes (Kim et al., 2019).

### Differential Gene Expression Analysis

Differential expression analysis was performed using the edgeR package (v3.42.4) in R/Bioconductor (Robinson et al., 2010). Raw counts were normalised using the trimmed mean of M-values (TMM) method. Genes with low expression (counts per million < 1 in fewer than two samples) were filtered out prior to statistical analysis. The differentially expressed genes between the treatments were analysed to identify the candidate genes involved in pathogen-induced stress in different tomato cultivars, and SlCBL1-2 (Solyc08g007160.3) was selected for further studies.

### Validation by quantitative real time polymerase chain reaction (qRT-PCR)

The total RNA isolation from the leaves of control and *A. solani* infected tolerant and susceptible tomato cultivar at 8 dpi was done using Monarch^®^ Spin RNA Isolation Kit (New England Biolab, USA) and the first strand cDNA was synthesized using PrimeScript™ RT Reagent Kit (Takara). The qRT-PCR analysis was performed in QuantStudio 5 (Applied Biosystems) using PowerUp™ SYBR™ Green Master Mix (ThermoFischer Scientific). The qRT-PCR data was normalised using Tubulin as the internal control. The primers used for qRT-PCR are provided in Table S1. The relative gene expression levels were calculated using the 2^-△△Ct^ method.

### Expression and Purification of SlCBL1-2 protein

The gene encoding canonical SlCBL1-2 was synthesised with optimal codon usage for expression in *E. coli* and subcloned into pET-28a (+) with 6x His tag on upstream and downstream region. SICBL1-2 protein was expressed and purified according to standard protocol (Cherukandy et al., 2023). SlCBL1-2-pET28a was transformed into *Escherichia coli* BL21(DE3) cells, and a single colony was grown in LB medium (HiMedia) containing kanamycin (50 mg mL^−1^) overnight at 150 rpm and 37 °C. The overnight culture was used to inoculate fresh LB medium and, once the culture reached an optical density at 600 nm (OD_600_) of 0.6, 1 mM isopropyl β-D-1-thiogalactopyranoside (IPTG) was added and the culture incubated at 37 °C for 4 h. The bacterial culture was centrifuged for 20 min at 4,000 × g and the resulting pellet was sonicated in buffer comprising 20 mM Tris pH 7.4, 200 mM NaCl and 5% (v/v) glycerol, followed by centrifugation at 10,000 × g for 20 min. The supernatant was collected and loaded onto a Ni–NTA affinity column connected to a peristaltic pump. Different concentrations of imidazole (50mM, 100mM, 200 mM, 400mM, 600mM, 800mM, 1M) were loaded on the column to elute our protein of interest. All the eluted fractions were subjected to SDS-PAGE (12.5%) and the imidazole from the pooled fractions was removed by washing with buffer containing 20 mM EDTA. The protein was concentrated using an Amicon^®^ Ultra Centrifugal Filter and further purified using gel filtration chromatography using Superdex 75 column attached to AKTA (GE Healthcare) FPLC system. The purified fractions were concentrated and estimated using a UV-visible spectrophotometer (using the Beer-lambert’s formula). Western blot procedure was performed according to the standard protocol using anti His antibody (GenScript, Cat.No.A00186) as the primary antibody (Towbin et al., 1979).

### Circular Dichroism (CD) Spectroscopy

Far (200−250 nm) and near-UV (250 – 320 nm) CD spectra of SlCBL1-2 were recorded using circular dichroism spectrophotometer (Chirascan CD spectrophotometer, United Kingdom) using a cuvette of path length of 0.2 cm. The spectra were recorded in the buffer containing 20 mM Tris and 50 mM NaCl at pH 7.4 both presence and absence of 1 mM CaCl_2_ at 25°C. The solvent baseline was corrected by subtracting the spectrum of the buffer alone. Each spectrum is an average of three replicates. The alpha helical content of SlCBL1-2 in the presence and absence of Ca^2+^ obtained from the CD spectra was analysed using BeStSel online tool (Micsonai et al., 2025).

### Fluorescence Spectroscopy

The fluorescence emission spectra of SlCBL1-2 protein were measured in FluoroMax Plus spectrofluorometer (HORIBA, Japan). The protein samples (2 μM) were dissolved in the buffer containing 20 mM Tris and 50 mM NaCl at pH 7.4, and the spectrum was recorded in the presence and absence of 1 mM CaCl_2_. The samples were excited at 295 nm, and the emission spectra were recorded from 310 to 400 nm with a slit width of 5 nm for both excitation and emission spectra. The buffer baseline spectrum was recorded using similar conditions and subtracted across all samples. Each spectrum is an average of three replicates.

### Isothermal Titration Calorimetry

Ca^2+^-binding to SlCBL1-2 protein and the associated thermodynamic properties were assessed using isothermal titration calorimeter (ITC) (Malvern MicroCal VP-ITC, United States) as described (Narayanasamy & Aradhyam, 2018). All the protein samples, buffer (20 mM Tris and 50 mM NaCl buffer at pH 7.4) and the ligand solution were degassed before titration. The cell containing 10 μM of SLCBL1-2 was titrated with 1mM CaCl_2_ at 25°C with a stirring speed of 750 rpm and a spacing of 150 s for 19 injections (2µL each). All the ITC data were subtracted from the background heat of dilution and the baseline adjustments were performed using the MicroCal ITC application supplied by the manufacturer. The stoichiometry (N), enthalpy (ΔH) and dissociation constant (K_D_) were obtained from the ITC data. The association constant (K_A_), variation in free energy (ΔG) and entropy (ΔS) were derived using the equations (Vallone et al., 2016), where K_A_ = 1/ K_D_, ΔG = -RT ln K_A,_ and ΔG = ΔH – TΔS.

### *In silico* and evolutionary analysis of SlCBL1-2 Protein

The gene and protein sequences of SlCBL1-2 (Solyc08g007160.3) were retrieved from Phytozome (https://phytozome-next.jgi.doe.gov/). The interacting partners of the SlCBL1-2 protein involved in signal transduction were analysed through the STRING (Search Tool for the Retrieval of Interacting Genes/Proteins). The SlCBL1-2 protein sequence was used as a query in NCBI BLASTP to retrieve the orthologous SlCBL1-2 from the Solanaceae members for understanding the EF-hand domain conservation and phylogenetic analysis. The sequences from other family species and the isoforms were removed, and we retrieved 19 sequences. The CBL1 protein sequences of *Arabidopsis thaliana* (thale cress), *Gossypium hirsutum* (cotton) and *Oryza sativa* (rice) were used as the outgroup. The multiple sequence alignment of all the sequences was aligned using MAFFT and visualised in Jalview (Waterhouse et al., 2009). The evolutionary relationships of these proteins were aligned using Multiple Sequence Comparison by Log-Expectation (MUSCLE) and the tree was constructed using the Maximum Likelihood (ML) method in the Molecular Evolutionary Genetics Analysis Version-11 (MEGA-11) software with 1000 bootstrap values. The tree was visualised and annotated using iTOL (Interactive Tree of Life)(Mackrill, 2024).

### Molecular Dynamic (MD) Simulation of SLCBL1-2

The EF-hands of SlCBL1-2 were analysed by comparing them with the classical canonical EF-hands (Gifford et al., 2007). Since the three-dimensional structure of SlCBL1-2 has not been determined, we have used the AlphaFold structure of SlCBL1-2 (AlphaFold DB:AF-A0A3Q7HHR6-F1) for our MD simulation studies. The structure was validated using the Ramachandran plot. For predicting and docking the Ca^2+^ in the binding site, MIB2 server was used (Lu et al., 2022). The MD simulation of the SlCBL1-2 in the presence and absence of Ca^2+^ was carried out using the GROningen MAchine for Chemical Simulations (GROMACS) simulation package (Hess et al., 2008). For apo and holo SlCBL1-2 protein, the CHARMM27 all-atom force field was used. The protein was solvated in the SPC water model, neutralised by Na+ ions and using the steepest descent algorithm, the complex was minimised for 5,000 steps. To maintain constant temperature (300 K) and pressure (1 bar), V-rescale thermostat (NVT ensemble) and Parrinello–Rahman algorithm (NPT simulation phase) were used, respectively. and the simulation was performed for 100 ns for SlCBL1-2 in the presence and absence of Ca^2+^ (Amrutha et al., 2024). The MD simulations for apo and holo SlCBL1-2 was performed in triplicate. The overall stability and flexibility of the SlCBL1-2 protein structures were analysed using the root mean square deviation (RMSD) and root mean square fluctuation (RMSF) of the backbone atoms for the initial and final structure throughout the simulation using the tools available in the GROMACS software. The simulated structures were visualised using PyMol.

### Statistical analysis

All data were presented as mean ± standard error (SE) from three independent experiments (n=3). Statistical significance was performed using two-way ANOVA followed by Tukey’s test and defined as * (p < 0.05), **(p <0.01), and *** (p < 0.001).

## Authors contribution

N.R. conceived and designed the study. S.P.S. and M.C.A. performed the experiments. N.S., P.K. and N.R. analysed the data. S.P.S., M.C.A. and M.B.S. contributed to methodology. S.P.S. contributed to Writing – original draft. N.S., P.K., M.M. and N.R. contributed to Writing –review and editing. All authors have read and approved the final version of this manuscript.

## Acknowledgements

The authors S.P.S and N.R. thank the Director & Head, Centre for Advanced Studies in Botany, University of Madras, for providing the laboratory facility. S.P.S acknowledge the financial assistance provided by CSIR during this work. The authors S.P.S and N.R. thank the Prof. G.N.Ramachandran Central Instrumentation Centre, University of Madras, for fluorescence spectroscopy analysis and the Department of Chemistry, Indian Institute of Technology Madras, Chennai, for helping with CD spectroscopy and ITC experiments.

