## Supplementary Files for "Molecular characterization of pathogen-induced SlCBL1-2, a calcium sensor, reveals the structural basis of calcium sensing in tomato"

**Graphical Model**

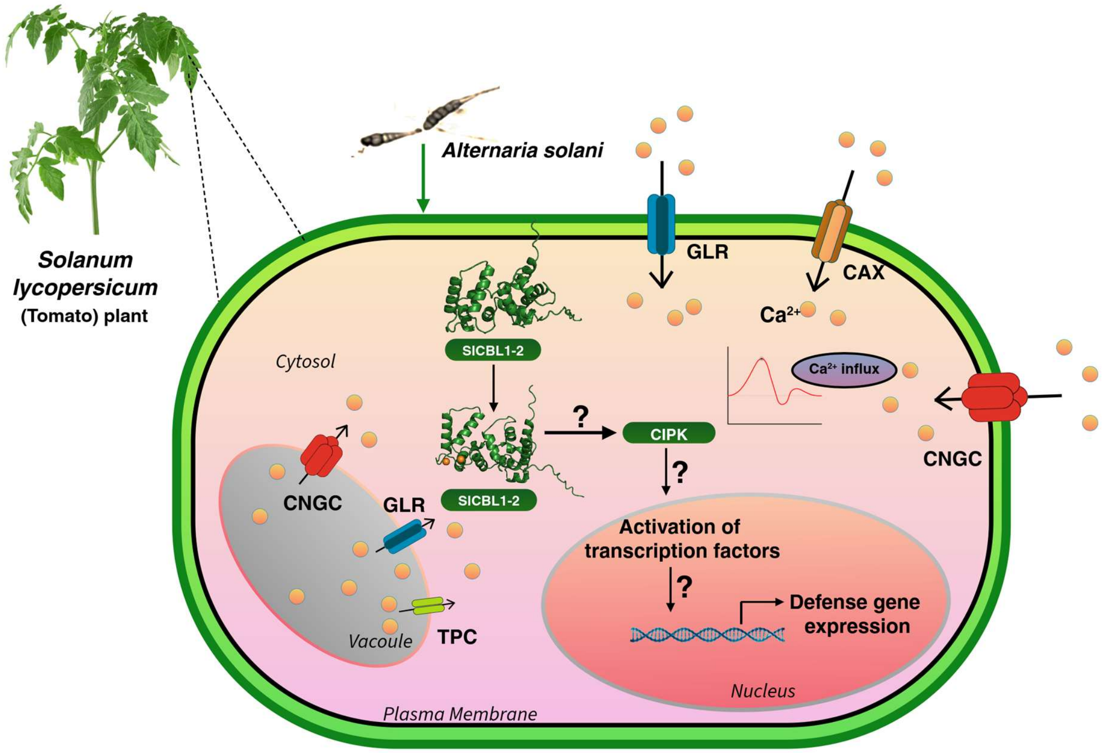


**Proposed model of the role of SlCBL1-2 in $Ca^{2+}$-mediated defense signalling in tomato against *Alternaria solani*.** *Alternaria solani*, a necrotrophic fungal pathogen, infects the tomato plant. In response to the fungal infection (biotic stress), $Ca^{2+}$-permeable channels localised to the plasma membrane, including cyclic nucleotide-gated channels (CNGCs), glutamate receptor-like channels (GLRs) and $Ca^{2+}$ /H+ exchangers (CAX), and $Ca^{2+}$ release from intracellular stores, including the vacuole, through two-pore channels (TPCs), activate cytosolic $Ca^{2+}$ signatures. Elevated cytosolic $Ca^{2+}$ is detected by a calcium sensor protein, SlCBL1-2, that undergoes conformational changes upon $Ca^{2+}$ binding. Upon activation, $Ca^{2+}$ bound to SlCBL1-2 interacts with CBL-interacting protein kinases (CIPKs) to generate a signalling complex that transduces calcium signals downstream. This signalling cascade activates transcription factors in the nucleus, resulting in the expression of defense-related genes responses against pathogen infection.

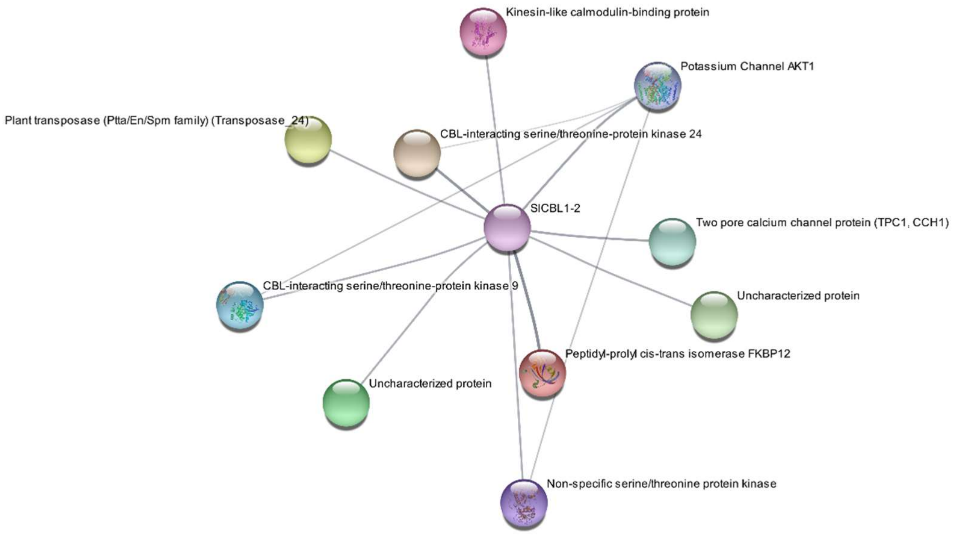


**Supplementary Figure 1. Protein-protein interaction network of SlCBL1-2.** The STRING based network reveals the predicted interactions of SlCBL1-2 with a few CBL-interacting protein kinases (CIPKs), the potassium channel AKT1, the two-pore calcium channel protein (TPC1/CCH1), and other proteins that might be involved in calcium signaling in *Solanum lycopersicum*.

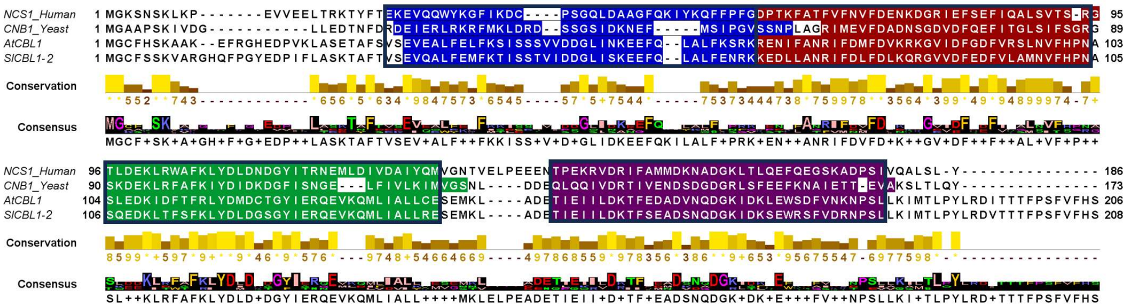

**Supplementary Figure 2.** Multiple sequence alignment of neuronal calcium sensor 1 in *Homo sapiens* (NCS1_Human), Calcineurin subunit B in *Saccharomyces cerevisiae* (CNB1_Yeast), CBL1 in *Arabidopsis thaliana* (AtCBL1), and CBL1-2 of *Solanum lycopersicum* (SlCBL1-2), highlighting the conserved EF-hand motifs. Blue colour represents EF-hand 1, red colour represents EF-hand 2, green colour represents EF-hand 3, and purple colour represents EF-hand 4.

**Table S1:** List of primers used for qRT-PCR.

| Name | Forward Primer (5'-3') | Reverse primer (5'-3') |
|---|---|---|
| SlCBL1-2 | ATGGGCTGTTTTAGCTCTAAG | TGAACATCTCAAATAAGGCCTG |
| Tubulin | GCGCCCTACCTACACCAATC | TGGATCCTGGGATAGGGGAC |

**Table S2:** Estimated secondary structure of apo and holo SlCBL1-2 using BeStSel.

| Secondary structure component | SlCBL1-2 (apo) (%) | SlCBL1-2 + $Ca^{2+}$ (%) |
|---|---|---|
| **Total helix** | **53.0** | **55.0** |
| Helix 1 (regular) | 33.2 | 34.6 |
| Helix 2 (distorted) | 19.8 | 20.4 |
| Antiparallel β-sheet | 0.0 | 0.0 |
| Parallel β-sheet | 0.0 | 0.0 |
| Turn | 10.8 | 12.7 |
| Others | 36.2 | 32.2 |
